# Structure-based identification and characterisation of novel inhibitors of K_Na_1.1 potassium channels, a stratified target for *KCNT1*-related epilepsy

**DOI:** 10.1101/779975

**Authors:** Bethan A. Cole, Rachel M. Johnson, Hattapark Dejakaisaya, Nadia Pilati, Colin W.G. Fishwick, Stephen P. Muench, Jonathan D. Lippiat

**Affiliations:** School of Biomedical Sciences, Faculty of Biological Sciences, University of Leeds, Leeds, LS2 9JT, U.K; Astbury Centre for Structural Molecular Biology, University of Leeds, Leeds, LS2 9JT, U.K; Autifony Srl, Istituto di Ricerca Pediatrica Citta’ della Speranza, Corso Stati Uniti, 4f, 35127, Padova, Italy; School of Chemistry, University of Leeds, Leeds, LS2 9JT, U.K

**Keywords:** potassium channel, KCNT1, K_Na_1.1, epilepsy, electrophysiology, molecular docking

## Abstract

Several types of drug-resistant epileptic encephalopathies of infancy have been associated with mutations in the *KCNT1* gene, which encodes the sodium-activated potassium channel subunit K_Na_1.1. These mutations are commonly gain-of-function, increasing channel activity, therefore inhibition by drugs is proposed as a stratified approach to treat disorders. To date, quinidine therapy has been trialled with several patients, but mostly with unsuccessful outcomes, which has been linked to its low potency and lack of specificity. Here we describe the use of a cryo-electron microscopy-derived K_Na_1.1 structure and mutational analysis to identify the quinidine biding site and identified novel inhibitors that target this site using computational methods. We describe six compounds that inhibit K_Na_1.1 channels with low- and sub-micromolar potencies, likely through binding in the intracellular pore vestibule. In preliminary hERG inhibition and cytotoxicity assays, two compounds showed little effect. These compounds may provide starting points for the development of novel pharmacophores for K_Na_1.1 inhibition, with the view to treating *KCNT1*-associated epilepsy and, with their potencies higher than quinidine, could become key tool compounds to further study this channel. Furthermore, this study illustrates the potential for utilising cryo-electron microscopy in ion channel drug discovery.

## Introduction

Gain-of-function mutations in the *KCNT1* gene are associated with severe, drug-resistant forms of childhood epilepsy (Barcia *et al*., 2012; Heron *et al*., 2012). Epilepsy of infancy with migrating focal seizures (EIMFS) and autosomal dominant nocturnal frontal lobe epilepsy (ADNFLE) were the first disorders found to be associated with *KCNT1*, with the more-recently described Ohtahara syndrome (Martin *et al*., 2014), West syndrome (Ohba *et al*., 2015), Lennox-Gastaut syndrome (Jia *et al*., 2019), and sleep-related hypermotor epilepsy (Cataldi *et al*., 2019; Rubboli *et al*., 2019). In addition to frequent seizures, patients may also present developmental delay, psychiatric and intellectual disabilities (Gertler *et al*., 1993; Lim *et al*., 2016). *KCNT1* encodes a sodium-activated potassium channel subunit, K_Na_1.1, which has previously been termed SLACK and Slo2.2 (Joiner *et al*., 1998; Yuan *et al*., 2003). Similar to other potassium channels, the functional proteins are formed by a tetramer of subunits, each of which possess six transmembrane alpha helices, a re-entrant pore loop between the fifth and sixth helix that forms the selectivity filter, and two intracellular regulation of conductance of potassium (RCK) domain (Hite *et al*., 2015). K_Na_1.1-containing channels are expressed throughout the central nervous system (Bhattacharjee *et al*., 2002; Rizzi *et al*., 2016) and are believed to have a stabilising effect on the membrane potential following sodium-influx during neuronal excitation (Liu & Stan Leung, 2004; Nanou *et al*., 2008; Budelli *et al*., 2009; Hage & Salkoff, 2012; Cervantes *et al*., 2013). Virtually all epilepsy-associated *KCNT1* mutations increase channel activity, though why epilepsy should arise is not understood, since potassium channel opening is usually associated with a decrease in neuronal excitability. One proposed mechanism, based on studies of human stem cell-derived neurons harbouring one such mutation, is that hyperexcitability can be caused by an enhanced sodium-dependent after-hyperpolarization, facilitating an increase in the rate of action potential firing (Quraishi *et al*., 2019).

Quinidine is a class I antiarrhythmic agent, which exerts its effects by non-selectively inhibiting cardiac cation channels at micromolar concentrations. Notably, quinidine also inhibits K_Na_1.1 channels (Yang *et al*., 2006), including those harbouring epilepsy-causing mutations, at similar concentrations, leading to the hypothesis that quinidine could reverse the gain-of-function and treat *KCNT1*-associated epilepsy (Milligan *et al*., 2014; Mikati *et al*., 2015). Limited improvement has been achieved in a small number of patients using quinidine therapy, but in the majority of cases it is ineffective (Mikati *et al*., 2015; Chong *et al*., 2016; Madaan *et al*., 2018; McTague *et al*., 2018; Mullen *et al*., 2018; Fitzgerald *et al*., 2019). The lack of selectivity and low potency of quinidine in inhibiting K_Na_1.1 channels, with IC_50_ values in the order of 0.1 mM (Yang *et al*., 2006; Rizzo *et al*., 2016), in the central nervous system without significantly affecting cardiac function is a concern and limits the dosing levels of quinidine (Mullen *et al*., 2018). Moreover, there is still a paucity of information on the binding site and the mode of action of K_Na_1.1 inhibitors. Other reported inhibitors of K_Na_1.1 are bepridil (Yang *et al*., 2006) and clofilium (de Los Angeles Tejada *et al*., 2012), both of which also have inhibitory effects on cardiac cation channels. The identification of alternative drugs that better target K_Na_1.1 are therefore desired (Milligan *et al*., 2014; Mikati *et al*., 2015).

With the development of direct electron detectors, more powerful microscopes, and improved data processing software, there has been an expansion of the use of cryo-electron microscopy (cryo-EM) and single particle analysis to determine high resolution structures of membrane proteins (Rawson *et al*., 2016). Bypassing the need to crystallise membrane protein samples makes this approach particularly attractive. One of the first high-resolution ion channel structures determined by cryo-EM was the chicken K_Na_1.1 channel, initially described at 4.5 Å resolution (Hite *et al*., 2015), with the inactive and activated conformations subsequently resolved to 4.3 and 3.8 Å, respectively (Hite & MacKinnon, 2017). The increase in resolution enables the generation of molecular models, though whether cryo-EM is presently able to deliver ion channel structural data that can be utilised in computer-aided drug discovery (CADD) or *in silico* drug design has not been fully explored.

In this study, we used the cryo-EM-derived structures of the K_Na_1.1 channel to model quinidine binding *in silico*. Having identified the intracellular pore vestibule as the likely site for known inhibitors to bind, we then conducted virtual high-throughput screening of a library of commercially-available compounds with the view to identify more potent and/or selective binders. We now report the identification of six diverse compounds that inhibit K_Na_1.1 more potently than the quinidine.

## Materials & Methods

### Model preparation and molecular docking

Models of the K_Na_1.1 pore were generated in UCSF Chimera using the atomic models of the putative closed and open tetrameric channel conformations (PDB:5U76 and 5U70, respectively (Hite & MacKinnon, 2017), and comprised residues 244 to 340 (S^244^AMF…LWME^334^) of each subunit. Automated docking was conducted using SwissDock (Grosdidier *et al*., 2011) and GLIDE (Friesner *et al*., 2004) in Maestro (Schrödinger). Virtual high-throughput screening was performed using GLIDE with the model of the channel in the putative open state and the Chembridge compound library consisting of 100,000 screening compounds were docked. Both protein and ligand-based docking approaches were used. Initially the compounds were screened in HTVS mode using the inner pore vestibule as the binding site, ranked according to their predicted binding affinities and the top-scoring 10,000 compounds were taken forward and subsequently docked using the higher precision SP mode. Approximately 100 top-scoring compounds were visually inspected using PyMOL to identify the binding interactions with the protein. For the ligand-based approach, ROCs (Hawkins *et al*., 2007) was used to overlay the Chembridge library of compounds over the predicted bepridil binding pose. The compounds which were predicted to overlay with bepridil were subsequently docked into the inner pore vestibule to determine the predicted binding affinities and were analysed to determine whether they would form any interactions to the protein. The structures of all of the best scoring compounds from both approaches were analysed to determine the ‘drug-like’ properties of the molecules. Of the top-scoring compounds which fit the criteria, 17 were then ordered from Chembridge (Chembridge Corp., San Diego, CA) and were shipped as dry stocks in 1 or 5 mg quantities before being dissolved in dimethylsulphoxide to stock concentration of 10 mM.

### Molecular biology and cell culture

A full-length K_Na_1.1 cDNA clone (IMAGE: 9054424) was obtained (Source Bioscience, Nottingham, U.K.) and the coding region subcloned into the pcDNA6-V5/His6 vector (Invitrogen). Mutations were introduced by the polymerase chain reaction using the New England BioLabs mutagenesis method and confirmed by sequencing (Genewiz, Takeley, U.K.). Due to the large size and high GC content of the insert, some mutations were generated in a plasmid containing the Bsu36I/BspEI (pore mutants) or SbfI/BsiWI (Y796H) restriction fragment and then subcloned into the corresponding sites in the pcDNA6-K_Na_1.1 construct. Human embryonic kidney (293) cells were cultured in Dulbecco’s modified Eagle medium (DMEM with GlutaMax, Invitrogen), supplemented with 10% v/v foetal bovine serum, 50 U/ml penicillin, and 0.05 mg/ml streptomycin. Cells were co-transfected with pcDNA6-K_Na_1.1 and pEYFP-N1 plasmid DNA using Mirus TransIT-X2 reagent (Geneflow, Lichfield, U.K.) and were plated onto borosilicate glass cover slips for electrophysiological experiments, which were conducted 2 to 4 days later.

### Electrophysiology

Unless otherwise stated, all chemicals were obtained from Sigma-Aldrich (Gillingham, U.K.). Patch pipettes were pulled from thin-walled borosilicate glass (Harvard Apparatus Ltd, Edenbridge, Kent, UK), polished, and gave resistances of 1.5 to 2.5 MΩ in the experimental solutions. The pipette solution contained, in mM, 100 K-Gluconate, 30 KCl, 10 Na-Gluconate, 29 Glucose, 5 EGTA and 10 HEPES, pH 7.3 with KOH and the bath solution contained, in mM, 140 NaCl, 1 CaCl_2_, 5 KCl, 29 Glucose, 10 HEPES and 1 MgCl_2_, pH 7.4 with NaOH. Currents were recorded from fluorescing cells at room temperature (20 to 22 °C) using the whole-cell patch clamp configuration using an EPC-10 amplifier (HEKA Electronics, Lambrecht, Germany), with >65 % series resistance compensation (where appropriate), 2.9 kHz low-pass filtering, and 10 kHz digitisation. For current-voltage analysis, cells were held at −80 mV and 400 ms pulses were applied to voltages between −100 and 80 mV. To evaluate inhibition by compounds, cells were held at −80 mV and 500 ms voltage ramps were applied from −100 to 40 mV at 0.2 Hz. Initially compounds, which were delivered by gravity perfusion, were applied serially at 10 μM (0.1 % final DMSO content) for 2 min, followed by at least 2 min wash with control solution before the next compound was added. Those compounds that that exhibited at least 40% current inhibition were analysed further by concentration-inhibition analysis: G/G_C_ = (1 + ([B]/IC_50_)^H^)^−1^ + c, where G is the conductance measured as the slope of the current evoked by the voltage ramp in the presence of the inhibitor, I_C_ is the control conductance in the absence of inhibitor, [B] is the concentration of the inhibitor, IC_50_ the concentration of inhibitor that yields 50% inhibition, H the slope factor, and c the residual current. For hERG currents, cells were held at −80 mV and 4 s depolarising pulses to +40 and then −50 mV were applied at 0.2 Hz. Data were analysed using Fitmaster (HEKA Electronics, Lambrecht, Germany), Microsoft Excel, and OriginPro 7.5 (OriginLab Corporation, Northampton, MA, USA). Data are presented as means ± s.e.m. (n = number of cells). Statistical analysis was carried out using SPSS (IBM analytics, Portsmouth, UK) with p<0.05 being considered significant.

### Cytotoxicity assay

Non-transfected HEK 293 cells were seeded at a density of 5×10^4^ cells/ well in a 96-well plate and incubated overnight at 37 °C in 5% CO_2_. Following exposure to three different concentrations of inhibitor for 24 hours, WST-1 reagent (Source Bioscience, Nottingham, U.K.) was added and cells were incubated for a further 2 hours. Inhibitor concentration ranges were selected to make comparison with effects on K_Na_1.1 currents. Cells were also treated with 10% v/v DMSO or 10 μg/ml blasticidin as positive controls. Absorbance at 450 nm (reference 650 nm) was measured using the Flexstation 3 microplate reader (Molecular Devices, Wokingham, UK). Cell viability was calculated as a percentage of the absorbance measured from untreated cells. Data were analysed using Microsoft Excel and OriginPro 7.5 (OriginLab Corporation, Northampton, MA, USA). Data are presented as mean ± s.e.m. (n = number of independent experiments). Statistical analysis was carried out using SPSS. Data were compared using an independent one-way ANOVA with Dunnett’s post-hoc test; p<0.05 was considered significant.

## Results

In the absence of potent and selective inhibitors of K_Na_1.1 potassium channels, we sought to use computational approaches to identify novel modulators. To identify a region in the K_Na_1.1 structure to focus *in silico* screening, we first sought to identify how compounds known to inhibit the channel exert their effects. Hypothesising that both quinidine and bepridil inhibit channels by occupying the inner pore vestibule we created a minimal structural model of the channel pore by removing the S1 to S4 and the cytosolic domains of the cryo-EM structures of the “closed” and “open” chicken K_Na_1.1 channel (Hite & MacKinnon, 2017). Using automated procedures, both inhibitors were docked into the pore in its closed conformation at two distinct sites: in the vicinity of F346 and M354 of the S6 segment of one subunit (Fig. 1A). In contrast, using the model of the open conformation, quinidine and bepridil could only be docked to the site involving F346 (Fig. 1A). To validate the docking, both residues (F346 and M354) were mutated to isoleucine and serine (Fig. 1B-C) and inhibition by quinidine and brepridil were evaluated further. Mutation of M354 had modest effects, with no significant effect on potency of quinidine, but the mutation of F346 to I346 reduced the potency approximately 10-fold (Fig. 2). We found that mutating these pore residues, particularly F346, also increased channel activity, with respect to current amplitude and a lower degree of rectification (Fig 1B-D). To exclude the possibility that this was not the cause of the reduced efficacy of the inhibitors, their effects on Y796H K_Na_1.1, an epilepsy-causing mutation that has similar effects, were explored. This gain-of-function epilepsy mutation (Heron *et al*., 2012) involves a site in the intracellular RCK domains distal to the transmembrane region, and so will not directly influence the binding of compounds to the pore. In contrast to the pore mutants, quinidine inhibited this mutant channel with a 3-fold increase in potency (Fig 2B).

**Figure 1:**
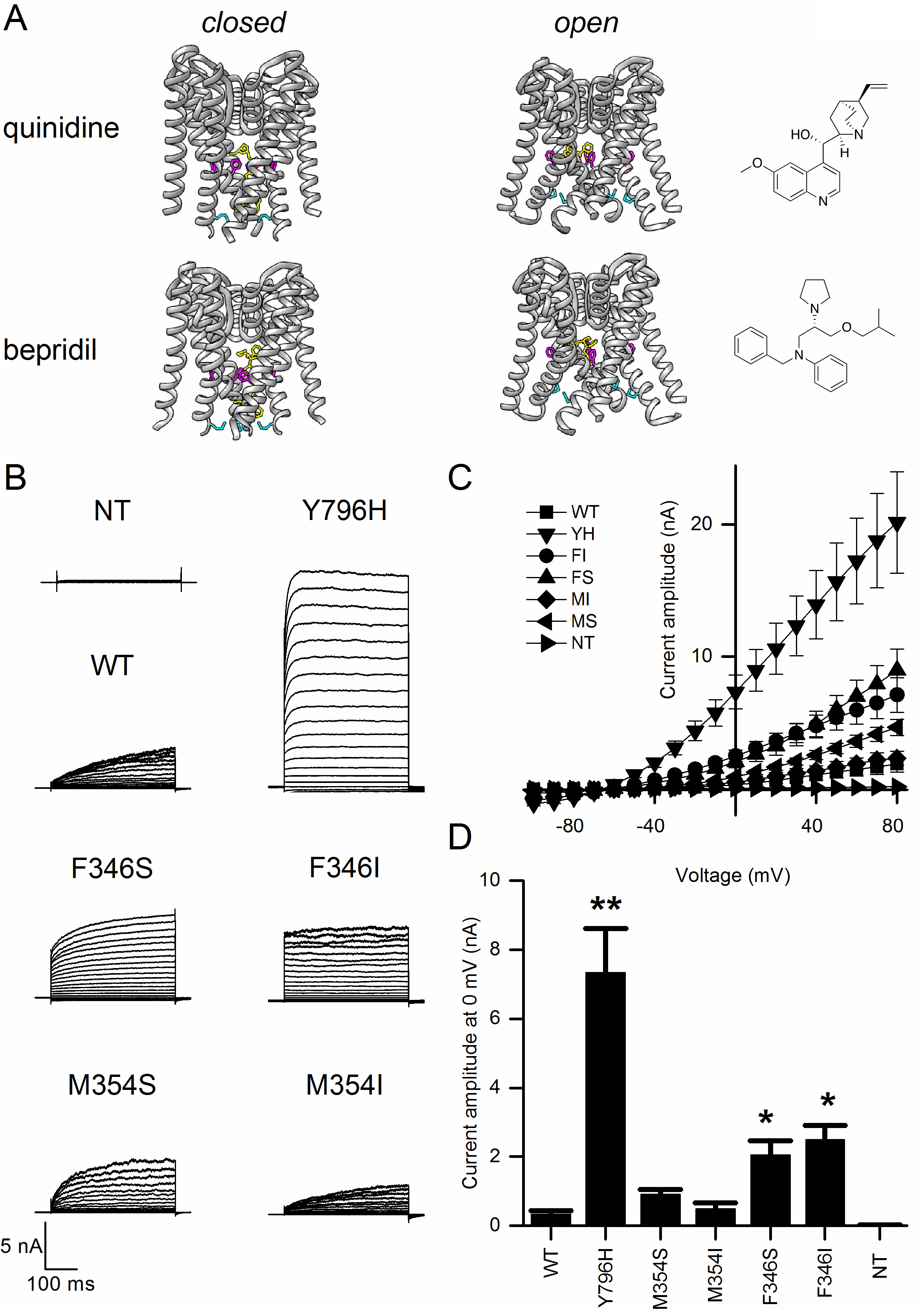
**A** Docking of quinidine and bepridil (yellow) into the KNa1.1 pore domain, comprising the S5, P-loop, and S6 of the closed and open conformational states (grey); side chains F346 (magenta) and M354 (cyan) are indicated. **B** Representative whole-cell currents and **C** mean (± s.e.m., n= 5 to 9 cells) current-voltage plots from non-transfected (NT) HEK 293 cells and cells transfected with wild-type (WT), pore mutant (F346S/FS, F346I/FI, M354S/MS, M354I/MI) or epilepsy-related mutant (Y796H/YH) KNa1.1. **D** Mean current amplitude at 0 mV from the data in **C**; *p<0.05, **p<0.005 compared to WT, independent one-way ANOVA with Games-Howell post-hoc test.

**Figure 2:**
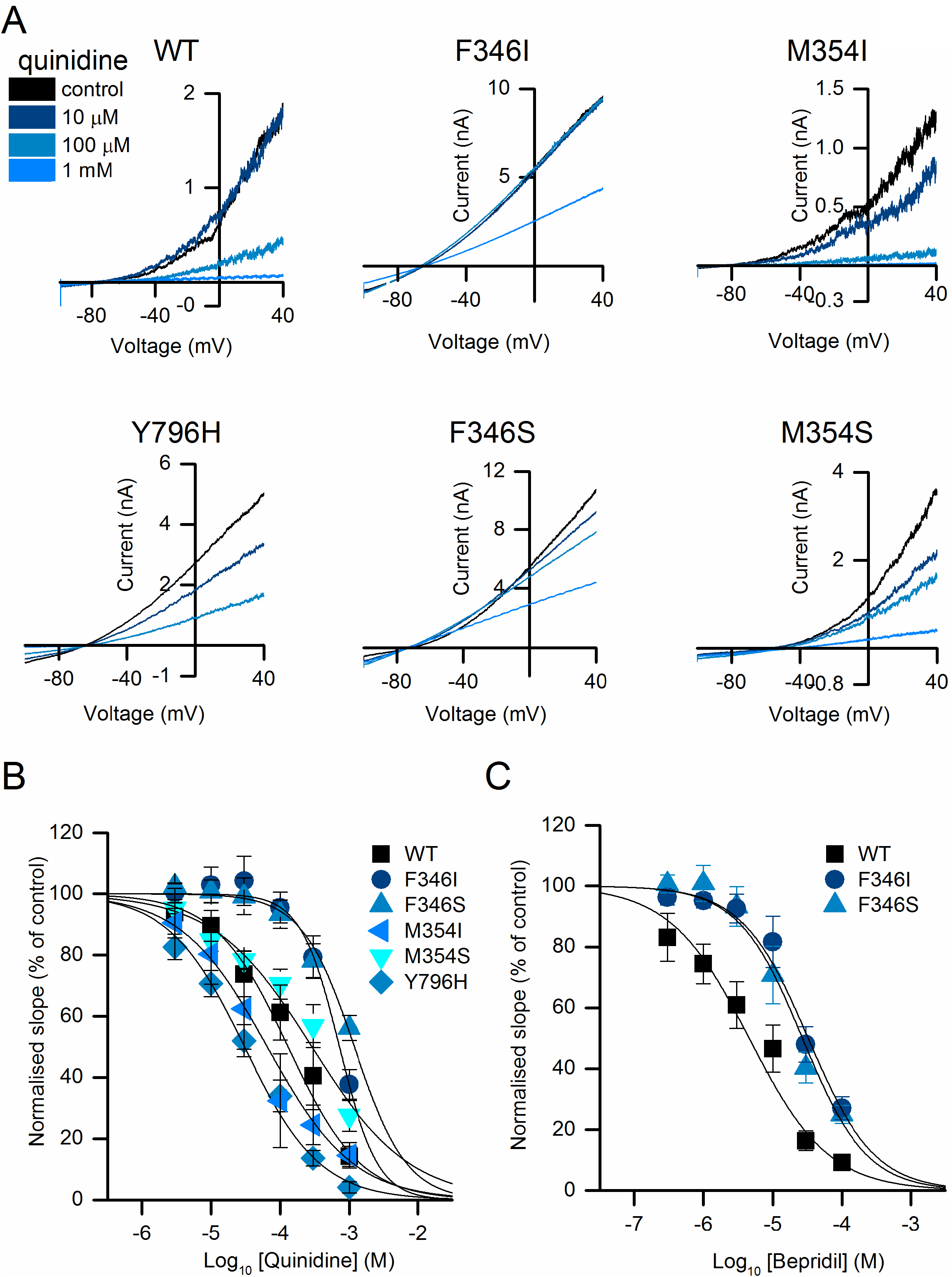
**A** Representative currents evoked by voltage ramps from cells expressing WT or mutant K_Na_1.1 with increasing concentrations of quinidine as indicated. **B** Concentration-inhibition plots for wildtype and mutant K_Na_1.1 channels in response to 3μM-1mM quinidine. IC_50_ for WT, 124.99 ± 34.52 μM (n=5 cells); F346I, 736.08 ± 94.09 μM (n=5); F346S, 1.23 ± 0.19 mM (n=4); M354I, 99.23 ± 49.61 (n=5); M354S, 247.16 ± 19.96 μM (n=5); Y796H, 38.00 ± 12.89 μM (n=5). **C** Concentration-inhibition plots for wildtype and mutant K_Na_1.1 channels in response to 0.3μM-100μM bepridil. IC_50_ for WT, 6.36 ± 2.12 μM (n=5); F346I, 35.91 ± 11.01μM (n=4); F346S, 23.43 ± 5.17 μM (n=5).

Having identified the internal vestibule, just below the selectivity filter, critical for inhibitor binding, we used this region in the minimal pore structure of the open channel conformation as the target for *in silico* compound screening. This approach was complemented by computational identification of compounds with structural similarity to bepridil, the most potent of the known inhibitors of K_Na_1.1. Both CADD techniques resulted in a list of compounds, ranked by their docking score and predicted binding affinities. A selection of 17 compounds (Supplementary Table 1), based on their ranking and commercial availability were then obtained and evaluated for their ability to inhibit human K_Na_1.1 channels expressed in HEK cells. At 10 μM, six of the compounds inhibited K_Na_1.1 currents by at least 40% and were selected for further analysis (Fig. 3A). Initially, and to validate inhibition in the inner pore vestibule, we tested the ability of these compounds to inhibit F346S K_Na_1.1, which had showed reduced sensitivity to both quinidine and bepridil. At 10 μM, the degree by which F346S K_Na_1.1 was inhibited by each compound was reduced, compared to WT KNa1.1 (Fig. 3A). Concentration-inhibition analysis (Fig. 3B-D) yielded mean IC_50_ concentrations ranging 0.6 to 7.4 μM with WT K_Na_1.1. In comparison, quinidine and bepridil inhibited WT K_Na_1.1 with IC_50_ values in the order of 125 μM and 6.4 μM, respectively (Fig. 2 and 3D). Inspection of the location of the inhibitors docked into the K_Na_1.1 pore domain indicated that each bind in the inner vestibule below the selectivity filter (Fig. 4). Binding appears to involve both hydrophobic interaction with S6 pore-lining residues and hydrogen-bonding with P-loop residues (Fig. 4 and Supplementary Table 1).

**Figure 3:**
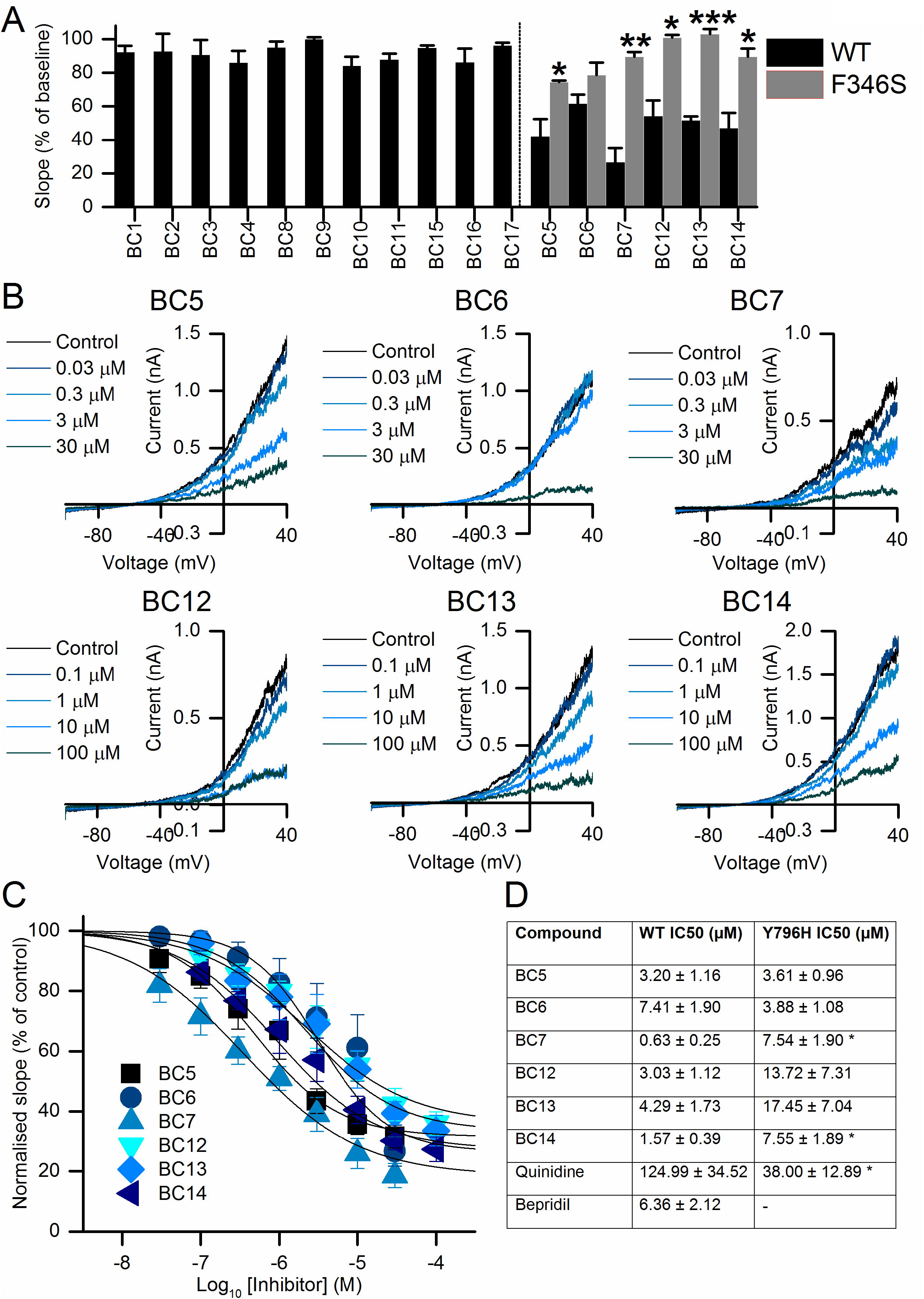
Functional evaluation of top-scoring molecules from *in silico* docking. **A** Mean (± s.e.m., n = 3-4 cells) WT K_Na_1.1 conductance measured as the slope of the current evoked by depolarising voltage ramps, relative to baseline, in the presence of 10 μM test compound; with those that were active (right of dashed line) counter-tested with F346S K_Na_1.1 pore mutant (*p<0.05, **p<0.005, ***p<0.0005, T-test.). **B** Representative traces and **C** mean (± s.e.m., n= 5 to 7) concentration-inhibition plots for active inhibitors. **D** Summary table with mean potencies of compounds in inhibiting WT and Y796H K_Na_1.1 (n-values the same as reported above, n=3 for other Y796H data). *p<0.05 vs potency with WT K_Na_1.1 (Student t-test).

**Figure 4:**
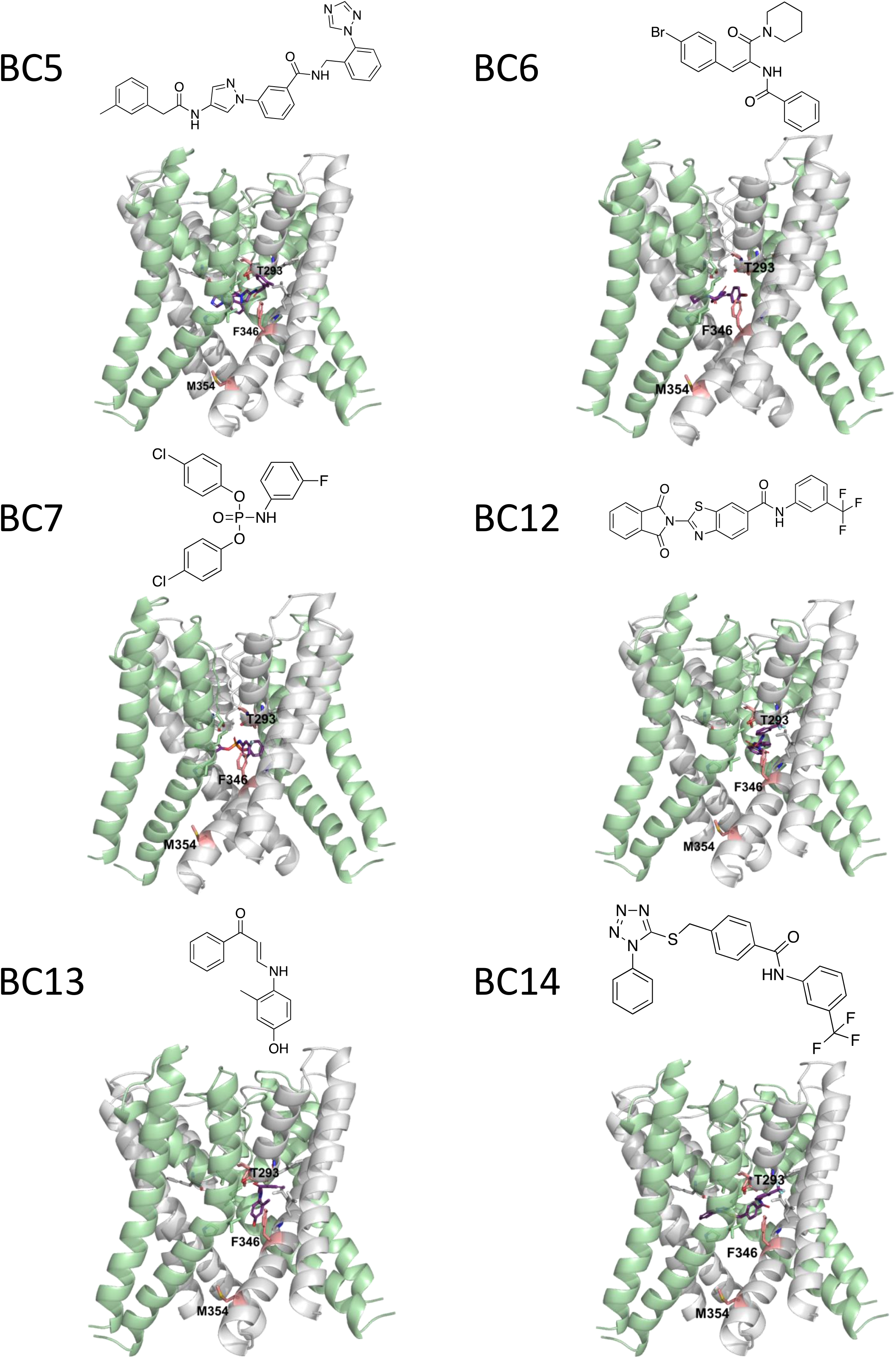
Chemical structures of inhibitors (magnenta) and their docked poses in the K_Na_1.1 pore domain. Side chains T293 in the P-loop, and F346 and M354 in the S6 segment, are indicated (pink).

The nature of the inhibition by these compounds was explored further. To be used therapeutically, K_Na_1.1 inhibitors would be required to inhibit channels that have epilepsy-causing amino acid substitutions. We therefore explored the potency of the inhibitors with Y796H K_Na_1.1. Each of the six compounds inhibited mutant channels with similar potency as with WT K_Na_1.1, though significantly lower potency was found with BC7 and BC14 (Fig. 3B and Supplementary Fig. 1). We also explored the importance of chemical groups in channel inhibition, using BC12 as an example, for which several analogues were commercially-available. Each analogue failed to reduce WT K_Na_1.1 channel currents by more than approximately 10% at 10 μM, and were deemed inactive (Supplementary Fig. 2).

Finally, to anticipate toxicological effects of these novel compounds, should they or derivatives be developed further, we studied their effects on hERG potassium currents and in a cellular toxicity assay. From measurements of tail current amplitudes, 10 μM BC5, BC6, and BC7 almost completely inhibited hERG channels expressed in HEK 293 cells (>80%, Fig. 5A, B). In contrast, BC14 had a partial effect (approx. 45%) at the same concentration, whilst BC12 and BC13 were even less effective, reducing currents by approximately 10-20%. In cytotoxicity assays, which involved exposing HEK 293 cells to compounds for 24h, only BC7 exhibited a concentration-dependent reduction in cell viability, at concentrations of 1 μM and above (Fig. 5C). With BC5, B6, and BC14, a reduction in cell viability was found at concentrations an order of magnitude higher than the IC_50_, whilst BC12 and BC14 had no effect at all concentrations tested, up to and including 100 μM (Fig. 5C). Blasticidin (10 μg/ml) and DMSO (10% v/v) reduced cell viability in the order of 45% and 90%, respectively.

**Figure 5:**
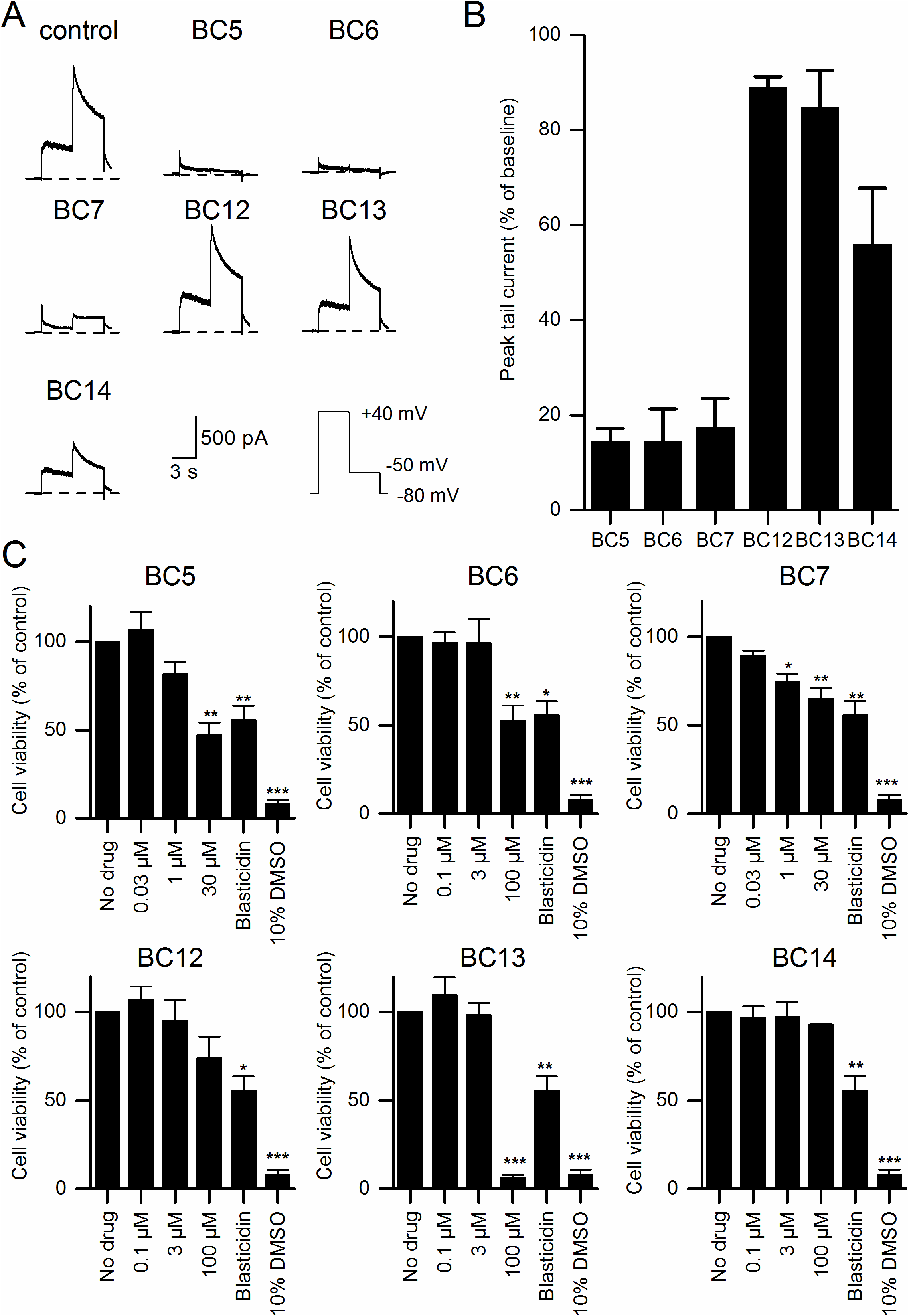
Preliminary toxicological data. **A** Representative hERG whole-cell currents recorded from a transfected HEK 293 cell in the absence (control) and presence of 10 μM inhibitor, as indicated. **B** Mean (± s.e.m., n=3) tail current at −50 mV remaining in the presence of each inhibitor. **C** Cytotoxicity assays indicating mean (± s.e.m., n=3) viability of HEK 293 cells using WST reagent following overnight exposure to inhibitor at the indicated concentrations, with 10 μM blasticidin and 10% v/v DMSO as positive controls; *p<0.05, **p<0.005, ***p<0.0005, independent one-way ANOVA with Dunnett’s post-hoc test.

## Discussion

Analysis of compounds identified as potential K_Na_1.1 inhibitors by *in silico* docking yielded six previously-unknown compounds, each more potent than quinidine, which has been trialled clinically as a stratified treatment for *KCNT1*-associated epilepsy. The novel inhibitors described here are diverse and there is no clear pharmacophore, though BC6 and BC7 are amongst the scaffolds with structural similarity to bepridil. We propose that each of these compounds, in addition to quinidine and bepridil, inhibit the channel by blocking the pore via the intracellular vestibule. This is indicated by the reduced efficacy of each compound with the F346S pore mutation, and which is likely to be independent of the increased channel activity caused by the mutation, since with quinidine there was no similar loss of efficacy with Y796H, an epilepsy-causing mutation distal from the pore. This mode of action requires the inhibiting compound to traverse the plasma membrane and enter the pore via the cytoplasm. It is noteworthy that each of the novel compounds tested that had a calculated logP (cLogP) value of 3.2 or less did not exhibit inhibition of K_Na_1.1, and this may have been as a result of poor membrane permeability. By using a model of the ion channel pore that is putatively in the open conformation, we had anticipated that the novel inhibitors would achieve similar or even higher potency with the Y796H mutant K_Na_1.1, but this was not always the case. It is possible that apparent increases in potency may be attributed to the high level of basal channel activity with the mutant channel and incomplete series resistance compensation, which would underestimate channel inhibition at the lower concentrations. The increase in potency of quinidine with Y796H, relative to WT K_Na_1.1, however, is consistent with previous studies of other gain-of-function mutations (Rizzo *et al*., 2016).

Whilst the cryo-EM structures indicate that activation by sodium involves an expansion of the intracellular pore vestibule (Hite & MacKinnon, 2017), functional experiments with this and the closely-related K_Na_1.2 and K_Ca_1.1 (BK_Ca_) channels point to the selectivity filter and proximal hydrophobic residues, rather than an S6 helix bundle, as the location of the channel gate (Garg *et al*., 2013; Suzuki *et al*., 2016; Giese *et al*., 2017; Jia *et al*., 2018). This means that the inhibitors described here block at the channel gate and this should be a mode of inhibition that is efficacious with virtually all clinical gain-of-function mutations, independent of the mechanism by which increased open probability if achieved, rather than an inhibitor that binds to modulatory sites. An exception, however, may be the F346L mutation, which appears to have effects analogous to the F346 mutations described here, and which has already been shown to be resistant to block by 300 μM quinidine in a *Xenopus* oocyte expression system (McTague *et al*., 2018), though the pharmacological effects of quinidine on channels containing a mixture of F346L and wild-type subunits has not been explored.

Notwithstanding our preliminary efforts to characterise the potential toxicity of these inhibitors by studying inhibition of hERG channel currents and the effects in cell viability assays, we are unable to make any statement regarding their selectivity or safety ahead of any *in vitro* or *in vivo* investigation. Since the computational approach was based on the inhibition by quinidine and bepridil, one might expect a similar range of ion channels to be inhibited by the compounds described here. However, the low level of inhibition of hERG at 10 μM by BC12 and BC13, suggests that across the compounds there may be varying levels of selectivity, which could be further improved by synthesis and analysis of derivative compounds and testing their effects on a range of different cation channels. We do note, however, the characteristics of pan-assay interference (PAINS) in compounds BC6, BC7, and BC13, due to the presence of the conjugated carbonyl group, meaning there may be non-specific effects in other functional assays. The lack of effects of the four analogues of BC12 tested demonstrate the potential for generating an inhibitor pharmacophore from this compound, thus these may provide starting points for the development of more potent inhibitors.

The generation of high-resolution structural data by cryo-electron microscopy and single particle analysis has revolutionised structural biology. Importantly, circumventing the need to crystallise samples means that membrane proteins are more amenable to analysis, and also that human proteins, rather than homologues from prokaryotes and lower eukaryotes, now feature more prominently. This means that for *in silico* analysis and drug discovery, target proteins very close to human, if not the human protein itself, can be utilised. The pore domain of the chicken K_Na_1.1 subunit, however, is virtually identical to that of the human homologue and we demonstrate its suitability for *in silico* docking experiments. Our study, in addition to identifying novel K_Na_1.1 inhibitors, provides an exemplification of the use of cryo-electron microscopy-generated membrane protein structural data in structure-based drug discovery.

## Acknowledgments

Supported by a BBSRC-CASE PhD studentship in conjunction with Autifony Therapeutics Ltd awarded to B.A.C., and a Wellcome Trust PhD studentship awarded to R.M.J.(109158/B/15/Z).

## References

Barcia G, Fleming MR, Deligniere A, Gazula VR, Brown MR, Langouet M, Chen H, Kronengold J, Abhyankar A, Cilio R, Nitschke P, Kaminska A, Boddaert N, Casanova JL, Desguerre I, Munnich A, Dulac O, Kaczmarek LK, Colleaux L & Nabbout R. (2012). De novo gain-of-function KCNT1 channel mutations cause malignant migrating partial seizures of infancy. Nat Genet 44, 1255–1259.

Bhattacharjee A, Gan L & Kaczmarek LK. (2002). Localization of the Slack potassium channel in the rat central nervous system. J Comp Neurol 454, 241–254.

Budelli G, Hage TA, Wei A, Rojas P, Jong YJ, O’Malley K & Salkoff L. (2009). Na+-activated K+ channels express a large delayed outward current in neurons during normal physiology. Nat Neurosci 12, 745–750.

Cataldi M, Nobili L, Zara F, Combi R, Prato G, Giacomini T, Capra V, De Marco P, Ferini-Strambi L & Mancardi MM. (2019). Migrating focal seizures in Autosomal Dominant Sleep-related Hypermotor Epilepsy with KCNT1 mutation. Seizure 67, 57–60.

Cervantes B, Vega R, Limon A & Soto E. (2013). Identity, expression and functional role of the sodium-activated potassium current in vestibular ganglion afferent neurons. Neuroscience 240, 163–175.

Chong PF, Nakamura R, Saitsu H, Matsumoto N & Kira R. (2016). Ineffective quinidine therapy in early onset epileptic encephalopathy with KCNT1 mutation. Ann Neurol 79, 502–503.

de Los Angeles Tejada M, Stolpe K, Meinild AK & Klaerke DA. (2012). Clofilium inhibits Slick and Slack potassium channels. Biologics 6, 465–470.

Fitzgerald MP, Fiannacca M, Smith DM, Gertler TS, Gunning B, Syrbe S, Verbeek N, Stamberger H, Weckhuysen S, Ceulemans B, Schoonjans AS, Rossi M, Demarquay G, Lesca G, Olofsson K, Koolen DA, Hornemann F, Baulac S, Rubboli G, Minks KQ, Lee B, Helbig I, Dlugos D, Moller RS & Bearden D. (2019). Treatment Responsiveness in KCNT1-Related Epilepsy. Neurotherapeutics.

Friesner RA, Banks JL, Murphy RB, Halgren TA, Klicic JJ, Mainz DT, Repasky MP, Knoll EH, Shelley M, Perry JK, Shaw DE, Francis P & Shenkin PS. (2004). Glide: a new approach for rapid, accurate docking and scoring. 1. Method and assessment of docking accuracy. J Med Chem 47, 1739–1749.

Garg P, Gardner A, Garg V & Sanguinetti MC. (2013). Structural basis of ion permeation gating in Slo2.1 K+ channels. J Gen Physiol 142, 523–542.

Gertler T, Bearden D, Bhattacharjee A & Carvill G. (1993). KCNT1-Related Epilepsy. In GeneReviews((R)), ed. Adam MP, Ardinger HH, Pagon RA, Wallace SE, Bean LJH, Stephens K & Amemiya A. Seattle (WA).

Giese MH, Gardner A, Hansen A & Sanguinetti MC. (2017). Molecular mechanisms of Slo2 K(+) channel closure. J Physiol 595, 2321–2336.

Grosdidier A, Zoete V & Michielin O. (2011). SwissDock, a protein-small molecule docking web service based on EADock DSS. Nucleic Acids Res 39, W270–277.

Hage TA & Salkoff L. (2012). Sodium-activated potassium channels are functionally coupled to persistent sodium currents. J Neurosci 32, 2714–2721.

Hawkins PC, Skillman AG & Nicholls A. (2007). Comparison of shape-matching and docking as virtual screening tools. J Med Chem 50, 74–82.

HeronSE, Smith KR, Bahlo M, Nobili L, Kahana E, Licchetta L, Oliver KL, Mazarib A, Afawi Z, Korczyn A, Plazzi G, Petrou S, Berkovic SF, Scheffer IE & Dibbens LM. (2012). Missense mutations in the sodium-gated potassium channel gene KCNT1 cause severe autosomal dominant nocturnal frontal lobe epilepsy. Nat Genet 44, 1188–1190.

Hite RK & MacKinnon R. (2017). Structural Titration of Slo2.2, a Na(+)-Dependent K(+) Channel. Cell 168, 390–399 e311.

Hite RK, Yuan P, Li Z, Hsuing Y, Walz T & MacKinnon R. (2015). Cryo-electron microscopy structure of the Slo2.2 Na(+)-activated K(+) channel. Nature 527, 198–203.

Jia Y, Lin Y, Li J, Li M, Zhang Y, Hou Y, Liu A, Zhang L, Li L, Xiang P, Ye J, Huang Z & Wang Y. (2019). Quinidine Therapy for Lennox-Gastaut Syndrome With KCNT1 Mutation. A Case Report and Literature Review. Front Neurol 10, 64.

Jia Z, Yazdani M, Zhang G, Cui J & Chen J. (2018). Hydrophobic gating in BK channels. Nat Commun 9, 3408.

Joiner WJ, Tang MD, Wang LY, Dworetzky SI, Boissard CG, Gan L, Gribkoff VK & Kaczmarek LK. (1998). Formation of intermediate-conductance calcium-activated potassium channels by interaction of Slack and Slo subunits. Nat Neurosci 1, 462–469.

Lim CX, Ricos MG, Dibbens LM & Heron SE. (2016). KCNT1 mutations in seizure disorders: the phenotypic spectrum and functional effects. J Med Genet 53, 217–225.

Liu X & Stan Leung L. (2004). Sodium-activated potassium conductance participates in the depolarizing afterpotential following a single action potential in rat hippocampal CA1 pyramidal cells. Brain Res 1023, 185–192.

Madaan P, Jauhari P, Gupta A, Chakrabarty B & Gulati S. (2018). A quinidine non responsive novel KCNT1 mutation in an Indian infant with epilepsy of infancy with migrating focal seizures. Brain Dev 40, 229–232.

Martin HC, Kim GE, Pagnamenta AT, Murakami Y, Carvill GL, Meyer E, Copley RR, Rimmer A, Barcia G, Fleming MR, Kronengold J, Brown MR, Hudspith KA, Broxholme J, Kanapin A, Cazier JB, Kinoshita T, Nabbout R, Consortium WGS, Bentley D, McVean G, Heavin S, Zaiwalla Z, McShane T, Mefford HC, Shears D, Stewart H, Kurian MA, Scheffer IE, Blair E, Donnelly P, Kaczmarek LK & Taylor JC. (2014). Clinical whole-genome sequencing in severe early-onset epilepsy reveals new genes and improves molecular diagnosis. Hum Mol Genet 23, 3200–3211.

McTague A, Nair U, Malhotra S, Meyer E, Trump N, Gazina EV, Papandreou A, Ngoh A, Ackermann S, Ambegaonkar G, Appleton R, Desurkar A, Eltze C, Kneen R, Kumar AV, Lascelles K, Montgomery T, Ramesh V, Samanta R, Scott RH, Tan J, Whitehouse W, Poduri A, Scheffer IE, Chong WKK, Cross JH, Topf M, Petrou S & Kurian MA. (2018). Clinical and molecular characterization of KCNT1-related severe early-onset epilepsy. Neurology 90, e55–e66.

Mikati MA, Jiang YH, Carboni M, Shashi V, Petrovski S, Spillmann R, Milligan CJ, Li M, Grefe A, McConkie A, Berkovic S, Scheffer I, Mullen S, Bonner M, Petrou S & Goldstein D. (2015). Quinidine in the treatment of KCNT1-positive epilepsies. Ann Neurol 78, 995–999.

Milligan CJ, Li M, Gazina EV, Heron SE, Nair U, Trager C, Reid CA, Venkat A, Younkin DP, Dlugos DJ, Petrovski S, Goldstein DB, Dibbens LM, Scheffer IE, Berkovic SF & Petrou S. (2014). KCNT1 gain of function in 2 epilepsy phenotypes is reversed by quinidine. Ann Neurol 75, 581–590.

Mullen SA, Carney PW, Roten A, Ching M, Lightfoot PA, Churilov L, Nair U, Li M, Berkovic SF, Petrou S & Scheffer IE. (2018). Precision therapy for epilepsy due to KCNT1 mutations: A randomized trial of oral quinidine. Neurology 90, e67–e72.

Nanou E, Kyriakatos A, Bhattacharjee A, Kaczmarek LK, Paratcha G & El Manira A. (2008). Na+-mediated coupling between AMPA receptors and KNa channels shapes synaptic transmission. Proc Natl Acad Sci USA 105, 20941–20946.

Ohba C, Kato M, Takahashi N, Osaka H, Shiihara T, Tohyama J, Nabatame S, Azuma J, Fujii Y, Hara M, Tsurusawa R, Inoue T, Ogata R, Watanabe Y, Togashi N, Kodera H, Nakashima M, Tsurusaki Y, Miyake N, Tanaka F, Saitsu H & Matsumoto N. (2015). De novo KCNT1 mutations in early-onset epileptic encephalopathy. Epilepsia 56, e121–128.

Quraishi IH, Stern S, Mangan KP, Zhang Y, Ali SR, Mercier MR, Marchetto MC, McLachlan MJ, Jones EM, Gage FH & Kaczmarek LK. (2019). An epilepsy-associated KCNT1 mutation enhances excitability of human iPSC-derived neurons by increasing Slack KNa currents. J Neurosci.

Rawson S, Davies S, Lippiat JD & Muench SP. (2016). The changing landscape of membrane protein structural biology through developments in electron microscopy. Mol Membr Biol 33, 12–22.

Rizzi S, Knaus HG & Schwarzer C. (2016). Differential distribution of the sodium-activated potassium channels slick and slack in mouse brain. J Comp Neurol 524, 2093–2116.

Rizzo F, Ambrosino P, Guacci A, Chetta M, Marchese G, Rocco T, Soldovieri MV, Manocchio L, Mosca I, Casara G, Vecchi M, Taglialatela M, Coppola G & Weisz A. (2016). Characterization of two de novoKCNT1 mutations in children with malignant migrating partial seizures in infancy. Mol Cell Neurosci 72, 54–63.

Rubboli G, Plazzi G, Picard F, Nobili L, Hirsch E, Chelly J, Prayson RA, Boutonnat J, Bramerio M, Kahane P, Dibbens LM, Gardella E, Baulac S & Moller RS. (2019). Mild malformations of cortical development in sleep-related hypermotor epilepsy due to KCNT1 mutations. Ann Clin Transl Neurol 6, 386–391.

Suzuki T, Hansen A & Sanguinetti MC. (2016). Hydrophobic interactions between the S5 segment and the pore helix stabilizes the closed state of Slo2.1 potassium channels. Biochim Biophys Acta 1858, 783–792.

Yang B, Gribkoff VK, Pan J, Damagnez V, Dworetzky SI, Boissard CG, Bhattacharjee A, Yan Y, Sigworth FJ & Kaczmarek LK. (2006). Pharmacological activation and inhibition of Slack (Slo2.2) channels. Neuropharmacology 51, 896–906.

Yuan A, Santi CM, Wei A, Wang ZW, Pollak K, Nonet M, Kaczmarek L, Crowder CM & Salkoff L. (2003). The sodium-activated potassium channel is encoded by a member of the Slo gene family. Neuron 37, 765–773.

