## Supplementary Table and Figures for "Structure-based identification and characterisation of novel inhibitors of K_Na_1.1 potassium channels, a stratified target for *KCNT1*-related epilepsy"

**Supplementary Material**


Bethan A. Cole[1], Rachel M. Johnson[1,2], Hattapark Dejakaisaya[1†], Nadia Pilati[3], Colin W.G. Fishwick[2,4], Stephen P. Muench[1,2], & Jonathan D. Lippiat[1]*.

[1]School of Biomedical Sciences, Faculty of Biological Sciences, University of Leeds, Leeds, LS2 9JT, U.K.

[2]Astbury Centre for Structural Molecular Biology, University of Leeds, Leeds, LS2 9JT, U.K.

[3]Autifony Srl, Istituto di Ricerca Pediatrica Citta' della Speranza, Corso Stati Uniti, 4f, 35127, Padova, Italy

[4]School of Chemistry, University of Leeds, Leeds, LS2 9JT, U.K.

[†]Present address: Department of Neuroscience, Monash University, Melbourne VIC 3004, Australia

**Supplementary Table 1:** Top-scoring compounds identified through virtual high-throughput screening and #bepridil overlay, assessed for $K_{Na}1.1$ inhibition at 10 μM. *Compounds that demonstrated >40% inhibition and were studied further.

| BC ID | PubChem ID | Chembridge ID | M.W. | cLogP | Docking score | Predicted H-bond |
|---|---|---|---|---|---|---|
| 1 | 42211371 | 10226003 | 433.5 | 2.1 | -8.731 | E326<br>T293 x2 |
| 2 | 42170356 | 18717381 | 450.5 | 2.611 | -8.841 | T392<br>S392 |
| 3 | 45198665 | 32500723 | 489.5 | 1.76 | -9.151 | T293 x2 |
| 4 | 72869676 | 38250229 | 360.4 | 2.9 | -8.88 | T293 x2 |
| 5* | 42481567 | 38627778 | 491.5 | 4.154 | -9.607 | T293<br>F291 |
| 6#* | 1283470 | 5143781 | 413.3 | 4.81 | -7.44 | T293 |
| 7#* | 2272824 | 5214461 | 412.2 | 7.05 | -7.189 | T293 x2 |
| 8# | 1376211 | 5422905 | 359.5 | 6.26 | -7.246 | n/a |
| 9# | 5341073 | 5574034 | 423.3 | 6.67 | -7.28 | T293 |
| 10 | 5342180 | 5690314 | 459.5 | 6.64 | -9.022 | F291<br>T293 x2 |
| 11 | 56902153 | 60526468 | 324.4 | 3.21 | -9.074 | F291 |
| 12* | 1330052 | 7040211 | 467.4 | 5.37 | -8.732 | F291<br>T293 x2 |
| 13* | 2185932 | 7364411 | 253.3 | 3.906 | -8.891 | F291 |
| 14* | 1243440 | 7942343 | 455.5 | 5.214 | -8.891 | n/a |
| 15 | 72922744 | 84082349 | 359.4 | 1.15 | -8.638 | F291<br>T293 x2 |
| 16 | 70780443 | 90931265 | 327.4 | 1.83 | -8.74 | F291 |
| 17 | 25249672 | 9273389 | 260.4 | 3.085 | -8.861 | F291 |

**Supplementary Figure 1**: Functional evaluation of novel inhibitors with $K_{Na}1.1$ channels harbouring the epilepsy-causing mutation Y796H. **A** Representative traces and **B** mean (± s.e.m., n= 3) concentration-inhibition plots for active inhibitors. **C** Summary table with mean potencies of compounds in inhibiting Y796H $K_{Na}1.1$; quinidine data from Figure 2.

**Supplementary Figure 2**: Evaluation of compounds closely-related to BC12. **A** Mean (± s.e.m., n = 3) $K_{Na}1.1$ conductance, measured as the slope of the current evoked by a depolarising voltage ramp in the presence of 10 μM test compound relative to control solution. **B** Chemical structures of compounds tested.

A

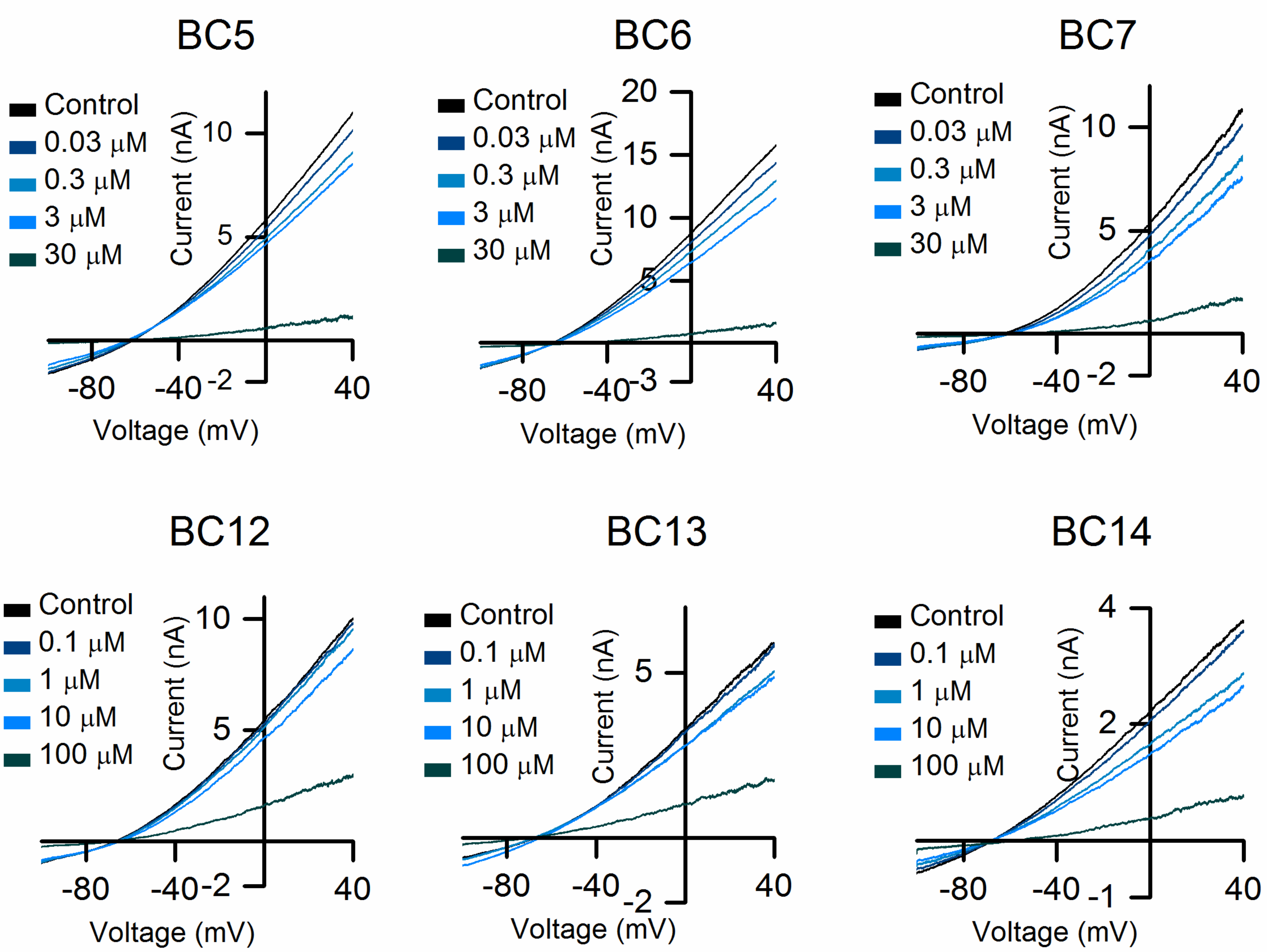


B

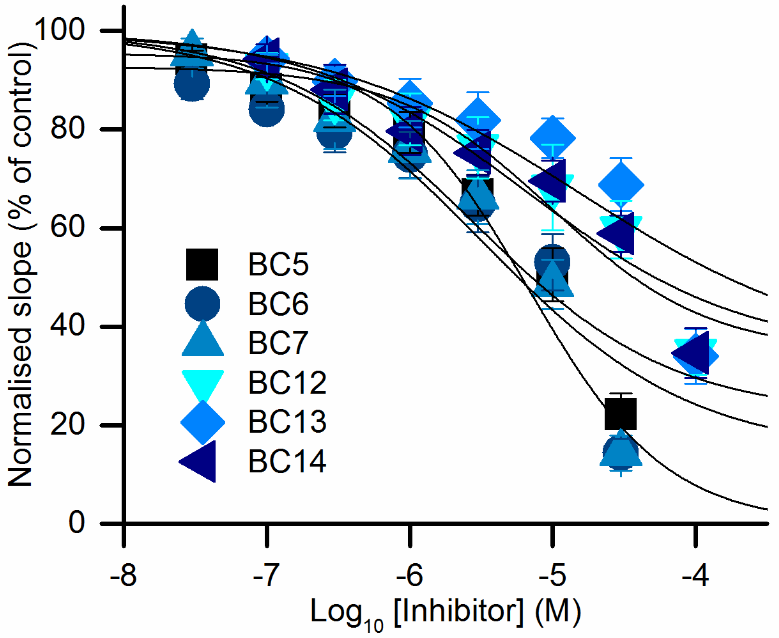


C

| Compound | IC50 (µM) |
|---|---|
| BC5 | 3.61 ± 0.96 |
| BC6 | 3.88 ± 1.08 |
| BC7 | 7.54 ± 1.90 |
| BC12 | 13.72 ± 7.31 |
| BC13 | 17.45 ± 7.04 |
| BC14 | 7.55 ± 1.89 |
| Quinidine | 38.00 ± 12.89 |

A

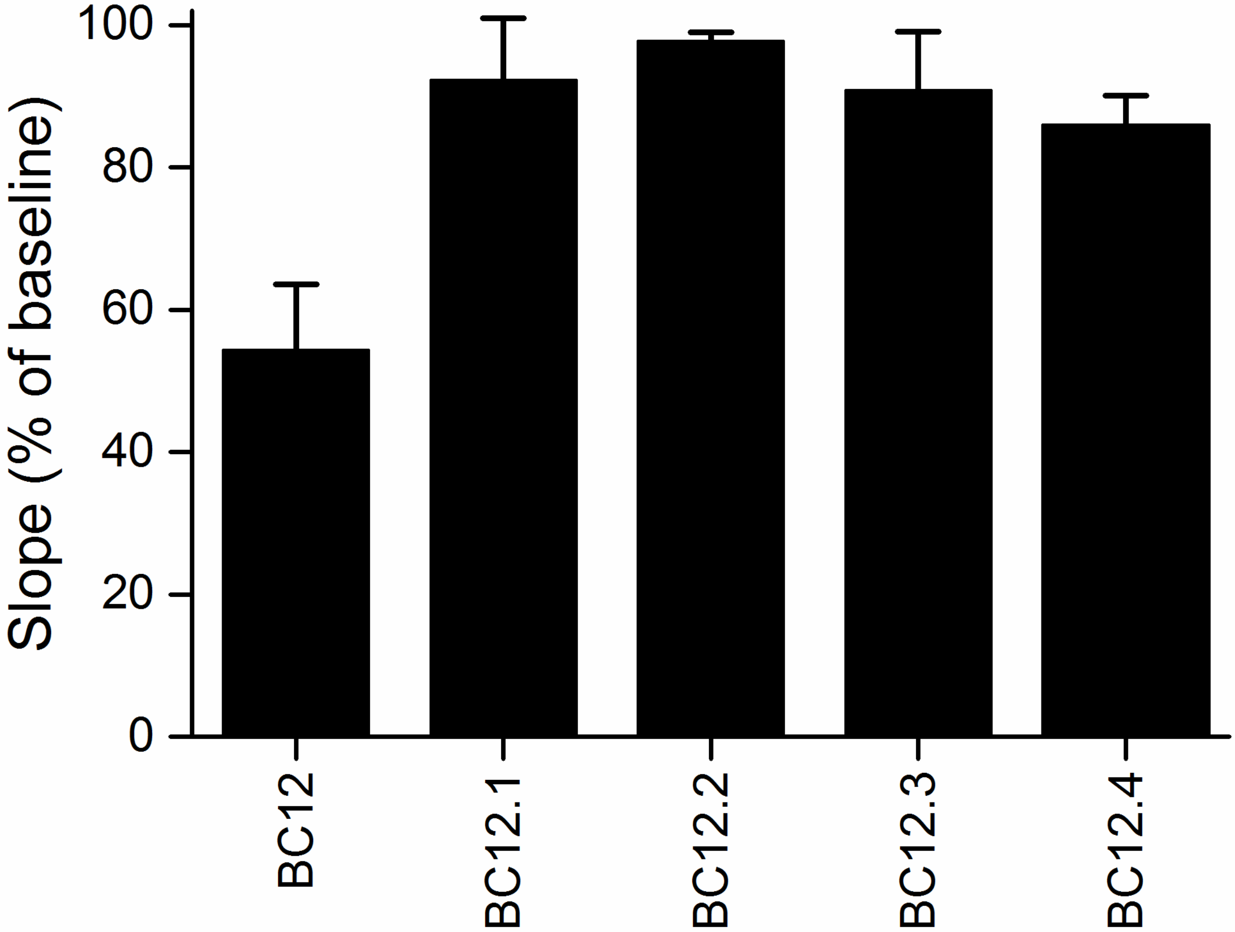


B

BC12

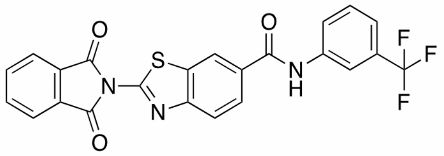

BC12.1

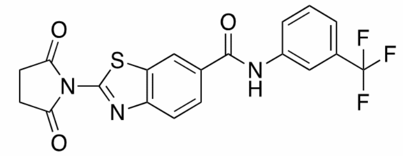

BC12.2

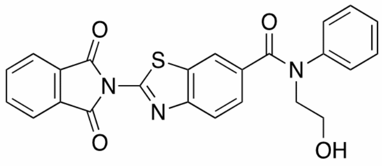

BC12.3

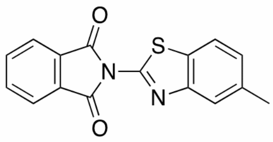

BC12.4

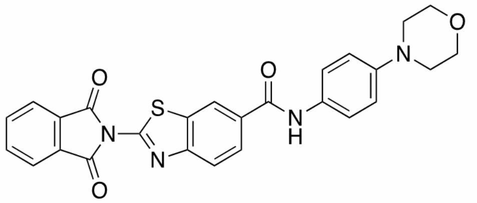
